# Clinically-guided mutation screening of two families with hereditary retinal disease

**DOI:** 10.1101/207068

**Authors:** Jian Li, Aierken Yiming, Ping Wang

## Abstract

Hereditary retinal disease (HRD) is a series of Mendelian diseases affecting the retina in the eye. The genetic basis of HRD is very complicated, with more than 100 disease-causing genes being identified. Though NGS has allowed rapid and large-scale mutation screening of Mendelian disease, the cost of NGS still prevents its universal application all over the world, for an accurate molecular diagnosis. Here, by clinical guidance from patient phenotypes, we performed targeted molecular diagnosis by direct Sanger sequencing of the most likely candidate gene in two families diagnosed with HRD. Then we identified two novel protein-truncating variants in the gene *CRB1*. Our results demonstrated the notion that molecular diagnosis and clinical diagnosis can be mutually supplemented and clinically guided direct sequencing is a cost-effective approach for molecular diagnosis and subsequent genetic counseling.

## Introduction

Hereditary retinal disease (HRD) is a group of human Mendelian disorders^1^. It includes a number of disease subtypes, including Leber congenital amaurosis, retinitis pigmentosa, cone-rod dystrophy, Stargardt disease, etc.. HRD usually features dysfunction and degeneration of photoreceptors in the retina. Photoreceptor is a highly specialized type of cell in the human body, and its survival and functions are controlled by a number of biological pathways. Therefore, mutations in more than one hundred genes can lead to HRD phenotype^2^. Currently, only 50%-70% of HRD cases can be solved by mutations in known HRD-associated genes. HRD is a relatively more common Mendelian disorder compared with other disease phenotypes, affecting around 1/2000 of the population^2^. This presents a particular need for improved molecular diagnosis.

The power of next-generation sequencing (NGS) has enabled large scale mutation discovery of human Mendelian disorders in recent years. It also speeds up the process of identifying novel disease-causing genes^3^. In the HRD field, the molecular diagnosis of disease cases also benefits a lot from a variety of NGS-based methods, including whole exome sequencing and targeted capture sequencing. Correspondingly, the molecular diagnosis rate of HRD cases has been significantly improved^4–13^, and a series of novel HRD-associated genes have been identified^14–33^.

However, the cost of NGS approach still prevents its universal application to all the populations affected by HRD, and direct Sanger sequencing is still a cost-effective way to screen for mutations in relatively small genes. Particularly, when a disease phenotype is very likely to be caused by mutations in one or two genes, it is probably not necessary to perform genome-wide mutation screening. Sanger sequencing demonstrates relatively short turn-around time for limited number of reactions, which is particularly useful in targeted analysis.

Here, by clinical evaluation of two disease families of HRD, we narrowed down the disease candidate to one gene, *CRB1*. Direct Sanger sequencing allowed us to identify two novel protein-truncating variants in this gene, thus showing the strength of Sanger sequencing in some specific scenarios.

## Methods and Materials

### Clinical diagnosis

The HRD disease families were recruited from Kashgar People’s Hospital, Kashgar, China for this study. Written informed consent was obtained from participating individuals. Specifically, we performed best visual acuity tests, color vision tests, fundus photography, visual field tests, fundus autofluorescence, and electroretinogram on all the patients for confirming the diagnosis. Family pedigrees were drawn based on interviews. Peripheral blood samples were collected from all available participating individuals. Genomic DNA was extracted from blood using the salting-out method^34^. This study adhered to the Declaration of Helsinki. All experimental methods were approved by the ethics committee of Kashgar People’s Hospital.

### Sanger sequencing and analysis

Primers were designed to amplify the coding exons of the candidate gene, *CRB1*. After PCR, the products were run on 1% agarose gel to confirm the identity. Then the products were sequenced on an ABI 3730xl machine. The sequence traces were analyzed on FinchTV to search for candidate mutations. GnomAD database was used to annotate the frequency of identified variants^35^. Variants with a population frequency more than 0.5% was excluded for additional analysis.

## Results

Two families with characteristic HRD phenotypes were included in this study. The Family 1 is a consanguineous family of Uyghur ethnicity. There are two affected individuals in the same generation (Figure 1A). The proband is a 36-year-old male. He had visual problems at the age of 6, starting from night blindness. Currently, he has a visual acuity of 20/80 for both eyes and the visual fields are restricted. ERG shows almost diminished responses of rod photoreceptors. Fundus images show bone spicules and very characteristic preserved para-arteriole retinal pigment epithelium (PPRPE) phenotype. This phenotype was shown in RP12 and this locus was later identified as the gene CRB1^36; 37^. The other affected individual in this family is a 23-year-old female. Similar to her brother, she had night blindness since 7 years old. Now her visual acuity is 20/50 for the left eye and 20/40 for the right eye. ERG test show about 50% reduction of rod responses. She also has the typical PPRPE phenotypes upon fundus photography. These two patients have five unaffected siblings.

**Figure 1A.**
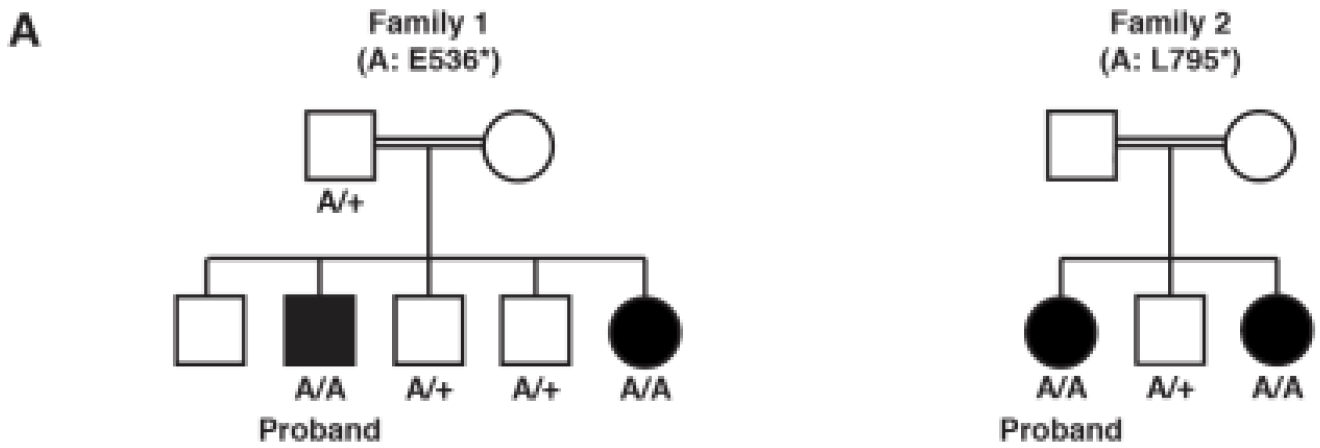
The pedigrees and genotypes of the two families in this study.

Family 2 is another consanguineous family also with Uyghur ethnicity. Two patients were identified in this family (Figure 1B). The proband is a 30-year-old female suffering from visual problems since childhood. She has the visual acuity 20/50 for both eyes and ERG responses are absent. She also has a sight nystagmus phenotype. Funduscopy revealed PPRPE phenotypes. She has an 23-year-old affected sister with typical RP phenotypes but without the PPRPE phenotypes.

**Figure 1B.**
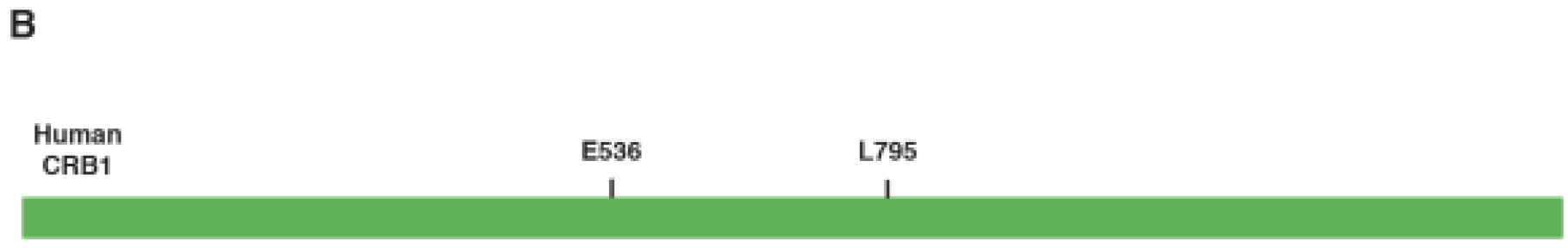
The location of the two novel *CRB1* variants identified in this study.

Due to the very typical phenotype of *CRB1*-related retinal disease presented in these two families, we decided to use direct Sanger sequencing for mutation screening. For both families, we chose the probands for direct sequencing. After Sanger sequence analysis, we identified two *CRB1* protein-truncating mutations in these two families. The proband in Family 1 has a homozygous stopgain mutation in exon 6 (NM_201253, c.1606G>T, p.E536*). Similarly, the proband in Family 2 also has a homozygous stopgain mutation in exon 7 (NM_201253, c.2384T>A, p.L795*). Both variants were not reported in HRD patients previously, and they are absent from the gnomAD control database, suggesting they are very rare. The two *CRB1* stopgain variants are not localized at the last exon, thus they are very likely to be loss-of-function alleles due to nonsense-mediated decay.

Finally, we performed additional Sanger sequencing for the two disease-causing mutations in the additional family members to test the genotype phenotype cosegregation. The genotypes are consistent with the recessive inheritance pattern (Figure 1A and B).

## Discussions

Sanger sequencing approach was developed several decades ago and it is a robust approach to identify germline variants existing in human DNA. Nowadays, with the availability of NGS, Sanger sequencing is still an indispensable validation method. In this study, we used the Sanger sequencing approach to successfully perform molecular diagnosis on two HRD cases based on their characteristic clinical presentations. These results showed that some typical clinical features of HRD are only caused by mutations in a small set of genes, and this information can help the researchers to narrow down the candidate genes for mutation screening.

CRB1 is a protein localized in the inner segment of the photoreceptor cells^38^. It is the orthologue of the Crumbs protein in the fly. CRB1 also has another paralogue, CRB2, also essential for photoreceptor maintenance^39^. However, it seems that CRB2 is indispensable in other tissue, thus mutations in *CRB2* lead to early-onset nephrotic syndromes^40^. Researchers have found that CRB1 is essential in maintaining the polarity and integrity of the photoreceptor cells. Mutations in *CRB1* also account for a significant proportion of HRD cases, especially for the characteristic PPRPE phenotype.

The mutations of some additional HRD-associated genes can also result in characteristic phenotypes, serving as candidate genes for direct Sanger sequencing when NGS-based approach is not readily available. For example, mutations in *ABCA4* can cause typical recessive Stargardt disease phenotypes^41^, and mutations in *CHM* can lead to choroideremia^42^. The utilization of the detailed gene-specific clinical outcomes would certainly improve the success rate of targeted Sanger sequencing.

The genetic etiology of HRD in the population of Uyghur ethnicity is still relatively understudied compared with some other populations, and one recent preliminary study investigated 12 HRD probands using NGS-based targeted exon capture sequencing^43^. Additional mutation screening efforts, especially by NGS-based approach, are needed to investigate the HRD molecular basis of population in this area. This will pave the way for a better genetic counseling and clinical outcome predictions for the HRD patients.

## Acknowledgements

This work was supported by Xinjiang Autonomous District Nature Science Funding (No.31760315). We thank Dr. Lei Wang for helpful suggestions on this project.

